# Excited State Relaxation Activation Energy (ESRAct) of Di-4-ANEPPDHQ Maps Nanoscale Molecular Organization in Biomembranes

**DOI:** 10.64898/2026.07.17.739057

**Authors:** Dishari Medda, Arpita Tripathy, Nirmalya Bag

## Abstract

Live cell plasma membranes show spatially heterogeneous liquid-ordered (Lo)-like and liquid-disordered (Ld)-like regions similar to the co-existing Lo/Ld phases observed in lipid vesicles. The Lo-like regions are relatively less hydrated and less polar due to tight packing of the membrane components compared to the Ld-like regions. The steady-state fluorescence spectra of Di-4-ANEPPDHQ (Di-4), a widely used polarity-sensitive probe, is blue or red shifted when solvated in less polar (Ld-like) or more polar (Lo-like) regions respectively. However, quantification of Di-4 fluorescence in blue and red channels for the evaluation of membrane phase state suffers from the lack of specific wavelength choice for these two channels and Di-4’s relatively higher concentration in Ld phase (red channel) due to its partitioning preference. To address these issues, we employed fluorescence lifetime of Di-4, a concentration independent photophysical parameter, to understand membrane biophysical properties. The fluorescence lifetime of Di-4 in lipid vesicles exhibits Arrhenius-like temperature dependence. Centred around this energetic feature of Di-4 photophysics, we developed a novel analytical module, namely excited state relaxation activation energy (ESRAct), that serves as an intrinsic descriptor of the membrane nano-environment sensed by this probe. We show that the ESRAact value scales with increasing disorder in nanoscale phase separation (i.e., ESRAct of pure Ld > mixed Ld/Lo > pure Lo phase). We then extended its applications to giant plasma membrane vesicles (GPMVs) isolated from MCF-7 cells and found that these vesicles exhibit nanoscale Lo/Ld co-existing phase within 16-37°C. We envisage wide applications of ESRAct to delineate plasma membrane phase behavior as well as general photophysical studies on other newly designed polarity-sensitive probes.

## INTRODUCTION

The plasma membranes (PM) of live cells serve as platform of cellular processes including the transmembrane signalling, host-pathogen interactions, cellular transport, cell-to-cell communications^1,2^. In recent times, it is becoming clear that membrane biophysical properties such as order and phase-like separation play crucial roles in these processes^1,3–14^. Quantitative measurements of membrane fluidity and order are therefore highly sought after. Among all methods, fluorescence based measurements are most widely used due to the sensitivity, compatibility with both non-living and living specimen^15–17^ as well as high resolution imaging techniques^18–20^, and recent advances in dye development^21,22^. Membrane fluidity and order are generally measured indirectly by monitoring changes in the fluorescence spectral properties of polarity-sensitive probes^23,24^. Briefly, tightly packed membranes are more ordered, which does not easily allow penetration of water molecules within plasma membranes^25,26^. This makes ordered membranes relatively more polar which is sensed by the polarity-sensitive probes such as Di-4-ANEPPDHQ (Di-4)^27^.

In general, plasma membrane is laterally heterogenous^13,18,28^ and is likely to possess co-existing liquid ordered (Lo) and liquid disordered (Ld) phases at nanoscale^5,29,30^. The Lo and Ld phases, which are experimentally observed in synthetic lipid vesicles^31–33^ and cell derived giant plasma membrane vesicles (GPMV)^34,35^, are respectively enriched in high-melting saturated and low-melting unsaturated or short chain saturated lipids^36^. Cholesterol plays a crucial role in lipid phase separation. It ‘fluidizes’ an otherwise tightly packed solid ordered (So) of gel phase lipid bilayer by breaking long-range order. In contrast, it induces short-range order when placed in relatively loosely packed liquid disordered (Ld) lipid bilayer^37–39^. These orthogonal effects of cholesterol on the packing of gel phase and Ld phase lipid essentially creates Lo phase. In essence, a mixture of high-melting and low melting lipids and cholesterol can show various types of phase states depending on the compositions^40^. Different phase states of the lipid membranes have differential levels of order and fluidity. Therefore, polarity-sensitive probes are very useful for investigations of membrane phase states and overall membrane order^15,24,41^.

A range of polarity-sensitive probes were developed over the last decades and they contributed enormously in our current understanding of membrane biophysical properties^6,18,42^ and their functional importance^13,43,44^. The steady-state emission spectra of these probes is blue and red shifted in polar and non-polar environments respectively. The sensitivity of local polarity however depends on the photophysical properties and location of a given probe^41^. Therefore all polarity-sensitive probes do not ‘sense’ the surrounding membrane environment equally^45,41^. Di-4-ANEPPDHQ (Di- 4) is among the most robustly used probe in this context. It is used to biophysical investigations in model membrane, live cell, live animal samples^15^. Given its suitable excitation wavelength^15^ for fluorescence microscopy, Di-4 labelled samples were imaged in confocal^46^, hyperspectral^47^, total internal reflection^48^, and super-resolution^20^ imaging modalities. Di-4 is a donor-π-acceptor type molecule (Fig. 1) which undergoes torsional rotation from the locally excited (LE) state to intramolecular charge transfer (ICT) state^45^. When Di-4 is localized in an environment having relatively tight lipid packing (less polar) its rotational dynamics is restricted. This also means that the ICT state of Di-4 in this environment is less stabilized (i.e., higher energy) than that in the more polar (and thus less ordered) environment. This is the underlying origin of the blue shift of Di-4’s emission spectrum in less polar environment. A recent report^49^ suggests that the sensor of Di-4 resides at the interface of lipid bilayer and surrounding water layer, rather than the lipid bilayer core. Therefore, Di-4 senses the hydration layer and hence the polarity at the bilayer surface which indirectly depends on the underlying lipid packing.

**Figure 1.**
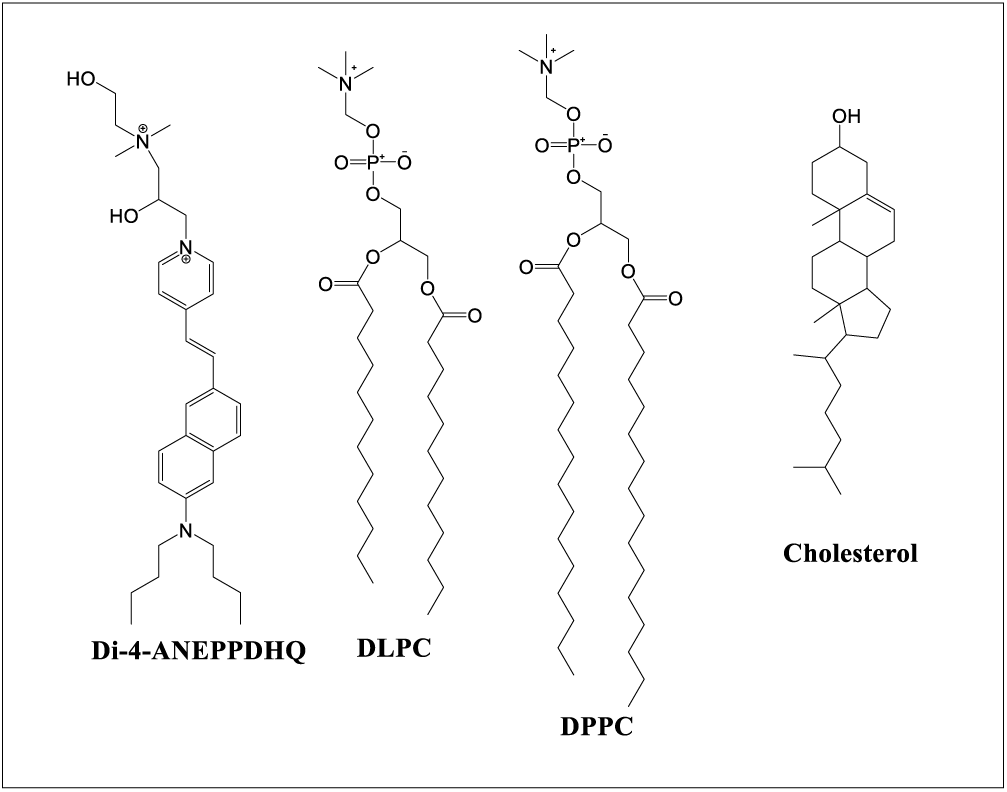
Chemical structures of the fluorescent probe, Di-4-ANEPPDHQ (Di-4), and lipids DLPC, DPPC, and Cholesterol, used in this study..

This steady-state spectral signature of Di-4 (and other polarity-sensitive probes) is used in the ratiometric analysis called ‘generalized polarization’ (GP) to delineate the changes of membrane order due to some perturbations^15^. The GP value is determined from fluorescence intensity of Di-4 at blue channel (less polar or more ordered) and a red channel (more polar and less ordered) are measured. The GP measurements were instrumental in deciphering the changes of membrane order in the biological processes^13,18,28,43,46,48^. This ratiometric GP analysis however suffers from two major concerns. First, the choice of wavelength is rather heuristic since the steady-state emission spectra of this probe in a phase separated ternary lipid bilayer (and more complex systems) does not show two obvious emission peaks for LE and ICT states correspondingly^47^. Second, the probe does not partition into ordered and disordered environments equally^27^ which results in differential probe concentration in these environments. Therefore, the intensity of the probe in a given emission channel is affected by both concentration and photophysics. This motivated us to employ concentration independent intrinsic properties of Di-4, namely fluorescence lifetime, to characterize polarity or fluidity of its local environment.

In this study, we systemically evaluate fluorescence lifetime of Di-4 at various compositions of ternary DLPC/DPPC/Cholesterol system^50^ using time correlated single photon counting (TCSPC) measurements. The melting temperature of DLPC and DPPC lipids is -2°C and 41°C respectively^51,52^. The phase diagram of DLPC/DPPC/Cholesterol system at room temperature was constructed using confocal imaging and Förster resonance energy transfer (FRET) spectroscopy on giant unilamellar vesicles^50^. Here, we measured temperature dependence of fluorescence lifetime of Di-4 which is doped in various large unilamellar vesicle (LUV) systems having compositions corresponding to Ld, So/Ld, Lo/Ld, and Lo phases of this ternary system based on the reported phase diagram^50^. We chose to use LUVs since their sizes are closer to biological vesicles and any phase separation in these systems will be nanoscopic which is also biologically relevant. We observed strong Arrhenius-like temperature dependence across all LUV systems. Centred around this energetics profile of Di-4 photophysics, we have developed an intrinsic descriptor, namely excited state relaxation activation energy (ESRAct), to get better insight on membrane nano-environments. The ESRAct values reflects the non-radiative transition activation energy between LE and ICT states. We discover that the ESRAct values of Di-4 follows the trend: Ld ∼ So/Ld > Lo/Ld > Lo and therefore distinctive ESRAct values of Di-4 can be tagged to underlying nanoscale phase separation. We finally estimated the ESRAct value in MCF-7 cell line derived GPMVs which resembles to that of Lo/Ld co-existing phase state. Overall, this study exploits complex excited state properties of Di-4 to develop ESRAct that provides quantitative information of nanoscale membrane biophysical properties.

## EXPERIMENTAL SECTION

### Materials

Lipids used in the study, 1,2-dilauroyl-sn-glycero-3-phosphocholine (DLPC) and 1,2- dipalmitoyl-sn-glycero-3-phosphocholine (DPPC) and cholesterol were bought from Avanti Polar Lipids (Sigma Aldrich, India). The membrane polarity sensitive probe Di-4 was purchased from Invitrogen. DL-Dithiothreitol (DTT) and Paraformaldehyde (PFA) were purchased from Sisco Research Laboratories Pvt. Ltd. (SRL, India) and Loba Chemie (India) respectively. Dulbecco’s Modified Eagle Medium (DMEM), fetal bovine serum (FBS), and Penicillin-Streptomycin (Pen-Strep) were purchased from ThermoFisher Scientific (India). 4-(2-hydroxyethyl)-1-piperazineethanesulfonic acid (HEPES) was bought from SRL, India. NaCl and CaCl_2_ were obtained from Sigma Aldrich, India. Stock solutions of lipids and cholesterol were prepared in chloroform. The Di-4 stock solution (0.75 mM) was prepared in dimethyl sulfoxide (DMSO) (Sigma Aldrich, India) and stored at 4°C.

### Preparation of cleaned round-bottomed flasks

The cleaning of glasswares were given elsewhere^53^. Briefly, 2 ml round-bottomed flasks were sonicated in a bath sonicator with10x diluted hellmanex reagent for 30 minutes at 25°C followed by vigorous rinsing with MilliQ water. The flasks were then again sonicated with 2M H_2_SO_4_ for 30 minutes. After washing, the flasks were subjected to sonication with MilliQ water for another 30 minutes. The cleaned flasks were stored in technical ethanol and air-dried after rinsing with chloroform before use.

### Preparation of Large Unilamellar Vesicles (LUVs)

We followed the LUV preparation protocols described previously^53^. To prepare LUVs, required amount of lipids were taken in clean round-bottomed flask and then lipid film was obtained by completely evaporating the solvent through rotary evaporator strictly for 3 hours. The obtained thin lipid film was hydrated in 2 ml aqueous buffer containing 10 mM HEPES and 150 mM NaCl (pH 7.4) for 20-30 minutes above the phase transition temperature of the highest meting lipids followed by vigorous vortexing for 1-2 minutes. The resulting milky lipid suspension of multilamellar lipid vesicles (MLVs) were immediately used or stored at 4°C overnight before the preparation of LUVs. On the day of measurement, the MLV suspension was sonicated in a bath sonicator (LMUC3D Model, LABMAN Scientific Instruments, India) above the chain melting temperature of lipids until translucent solution was obtained, thereby forming LUVs. After the formation of LUVs, Di-4 dye was added at a final concentration of 400 nM (an appropriate volume of stock solution (0.75 mM) directly added to LUV solution) and incubated for 20 minutes at room temperature and passed through 0.22 μm filter before measurements were taken. In the present study five different compositions of LUVs were prepared: as DLPC, DLPC/DPPC (1:1), DLPC/DPPC/CHOL (1:1 lipid mixture with 12 mol% cholesterol), DLPC/DPPC/CHOL (1:1 lipid mixture with 50 mol% cholesterol) and DPPC/CHOL (7:3). Total lipid concentration in 2 mL vesicle solution thus prepared was kept 1 mM.

### Cell culture and preparation of giant plasma membrane vesicles (GPMVs)

MCF-7 cells (kind gift from Prof. Mahitosh Mandal, Indian Institute of Technology, Kharagpur) were cultured in DMEM supplemented with 10% FBS and 1% penicillin and streptomycin. Cells were plated in T-25 flask before the isolation of GPMVs. They were isolated from MCF-7 cells using PFA/DTT method^35,54^. The confluency of the cells was 70-80% during the isolation of vesicles. Cells were carefully washed with vesiculation buffer (10 mM HEPES, 2 mM CaCl_2_, 150 mM NaCl) for two times followed by addition of 2 ml 25 mM PFA/ 2 mM DTT prepared in the same buffer. Cells were incubated for 4 hours at 37 °C and 5% CO_2_ environment to get sufficient GPMVs. After incubation suspension of GPMVs was taken in a centrifuge tube and stained with 400 nM Di-4 by keeping it for 20 minutes at room temperature. Preparations of vesiculation buffer was done in MilliQ water (passed through 0.22 μm filter).

### Steady-state Fluorescence Spectroscopy measurements

The fluorescence emission spectra of all the LUVs at different temperatures from 10°C to 46°C (±1°C) at 5°C interval were recorded using a spectrofluorimeter (F-7000, Hitachi, Japan) at scan speed 1200 nm/min. A refrigerated circulating water bath was used to control the temperature of the sample chamber. The samples were kept for 5 minutes at each temperature to equilibrate. Excitation wavelength (*λ_ex_*) was set at 476 nm and the emissions were collected from 496 to 700 nm wavelength window. The excitation and emission slit width of the monochromator was set at 5 nm.

### Time-resolved Fluorescence Spectroscopy measurement and data analysis

Di-4 labelled LUV or GPMV Sample (2 mL) was taken in a quartz cuvette and its fluorescence lifetime was measured. The time-resolved emission spectra at emission wavelength (*λ_em_*) 560 nm and 620 nm were recorded in reverse mode using DeltaFlex time correlated single photon counting (TCSPC) instrument (Horiba, Japan). A picosecond diode-pulsed laser at 476 nm with 20 MHz repetition rate was used as light source and the decays were taken under magic angle (54.7°) polarization using picosecond photon detector (PPD). A 515 nm long pass filter in emission pathway was used to eliminate scattering from the sample. The peak count value at each wavelength was 10,000 and the time window of the measurement was kept at 50 ns. The Instrument response function was measured using corresponding buffer solution. The temperature of the sample chamber was controlled using a Peltier cooling controller and the samples were equilibrated for 5 minutes at each temperature before data acquisition. The recorded time-resolved decays were analysed using EzTime decay analysis software (Horiba, Japan). The decays obtained in the LUVs were multi-exponential and they were fitted with bi-exponential decay curve following the mathematical equation:

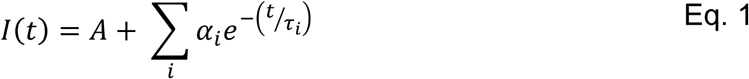

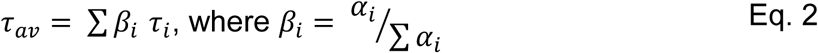

In the above equations, *I*(*t*) refers to the fluorescence intensity taken at time *t*, *A* denotes to the baseline correction, and *τ_i_* and *α_ι_* are fluorescence decay time of the *i* component of the emitted species and associated pre-exponential factor at time zero respectively. The goodness of the fit of the decays was judged by reduced chi-square values and random residual distribution. The amplitude-weighted average lifetime (*τ_av_*) is calculated from the lifetime values and contribution of all components.

We generally fit the time-resolved data with two-components (i.e., *i* = 2 is equation 1). These two fitted lifetimes were designated as *τ_LP_* and *τ_MP_* (in ns unit) corresponding to less polar (LP) and more polar (MP) regions of the LUVs (see Results and Discussions section for detail). Fluorescence lifetime is inversely related to excited state relaxation rate constant (*k*) and, thus for the LP and MP regions, the relaxation rate constants are given by 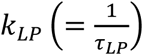 and 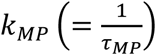 respectively. The temperature dependence of this rate constants follows Arrhenius equation (See Results and Discussion) as follows:

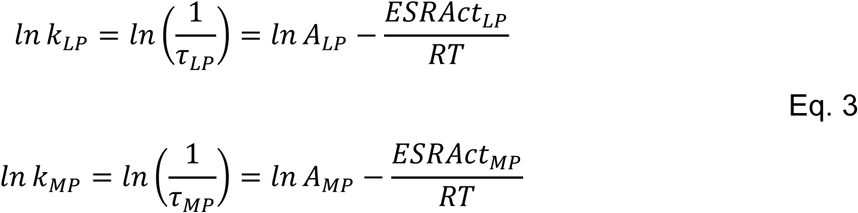

In the above equation, *A_LP_* and *A_MP_* are pre-exponential factor respectively for LP and MP regions. *ESRAct_LP_* and *ESRAct_MP_* are excited state relaxation activation energy of the probe on LP and MP regions. *R* is universal gas constant and *T* is absolute temperature.

### Laser Scanning Confocal Imaging

GPMVs were isolated from MCF-7 cells using above mentioned method followed by labelling with Di-4. Di-4 labelled GPMVs were placed in a custom-built glass-bottomed imaging petri dish and allowed to set for 15 minutes before starting imaging.

Confocal imaging of labelled GPMVs was performed using a commercial Olympus FV3000 inverted microscope equipped with a water immersion objective (60×magnification and 1.2 NA), Olympus) and GaAsP detector. Measurements were conducted at room temperature (298K). The samples were excited with a 488 nm laser line and the emission was detected at 560-660 nm wavelength range. The images were saved as .tiff file and further analysed in FIJI/ImageJ to determine the size of GPMVs.

## RESULTS AND DISCUSSION

The ratiometric GP measurements of Di-4 has long been used to investigate biophysical changes of plasma membranes in cellular processes^15^. The results after often linked to phase separation of the membrane. However, the steady-state fluorescence of this probe depends on the excited state photophysical properties which involve excited state torsional rotation enabling transition between LE to ICT state. The stabilization of the ICT state relative to the LE state depends on the hydration or polarity of the surrounding solvation layer. Therefore, we aimed to investigate the photophysical properties of Di-4 in biomimetic model lipid bilayers with well-defined phase diagram and links the excited state properties with underlying phase separation.

In this study, we evaluate fluorescence lifetime of Di-4 and its temperature dependence in single, two, and three-component LUVs comprising of DLPC, DPPC, and Cholesterol (Table 1) exhibiting various phase states^50^. At room temperature, DLPC and DPPC are respectively at fluid (*F*) or liquid disordered (Ld) and gel (*G*) or solid ordered (So) states. The bilayer systems are categorized as (Table 1): i) *F* - single component DLPC exhibiting Ld phase, ii) *FG* - two component DLPC/DPPC (1:1) exhibiting Ld/So co-existing phase, iii) *GC30* - DPPC/Cholesterol (7:3) exhibiting Lo phase, iv) *FGC12* - 1:1 DLPC/DPPC + 12 mol% cholesterol exhibiting Ld/Lo co- existing phase, and v) *FGC50* - 1:1 DLPC/DPPC + 50 mol% cholesterol exhibiting Lo phase. We specifically chose this ternary system because: a) The phase diagram of this system is well-established^50^, and b) In Lo-Ld co-existence composition, the system only shows nanoscopic phase separation^50^, rather than formation of microdomains, which is more relevant in cellular context^55–57^.

**Table 1:** Lipid compositions of different LUVs and their reported phase separation^50^.

| Bilayer Type | Composition |  |  | Phase<sup>50</sup> |
| --- | --- | --- | --- | --- |
|  | DLPC | DPPC | Cholesterol |  |
| F | 100 | 0 | 0 | Ld |
| FG | 50 | 50 | 0 | So/Ld |
| GC30 | 0 | 70 | 30 | Lo |
| FGC12 | 44 | 44 | 12 | Lo/Ld |
| FGC50 | 25 | 25 | 50 | Lo |

### Steady-state emission spectra of Di-4 labelled LUVs show weak temperature and moderate cholesterol dependence

Di-4 is weakly fluorescent in aqueous environment while its fluorescence significantly increases when embedded in hydrophobic assembly, for example, micelles (Fig. S1), lipid bilayers, and cell membranes. In agreement with the literature^58,59^, the emission spectra of Di-4 (Fig. 2) shows blue shift as the cholesterol concentration increases. Since cholesterol is known to increase local order in an otherwise fluid bilayer, the changes of Di-4’s emission spectra are often considered to mirror underlying changes of membrane order which is the primary distinguishing physical feature across various membrane phase states^27,59^. In the presence of cholesterol, the ICT state is less stabilized than in the Ld bilayer (*F* type) in which a larger amount of water molecules is allowed at the bilayer surface.

**Figure 2.**
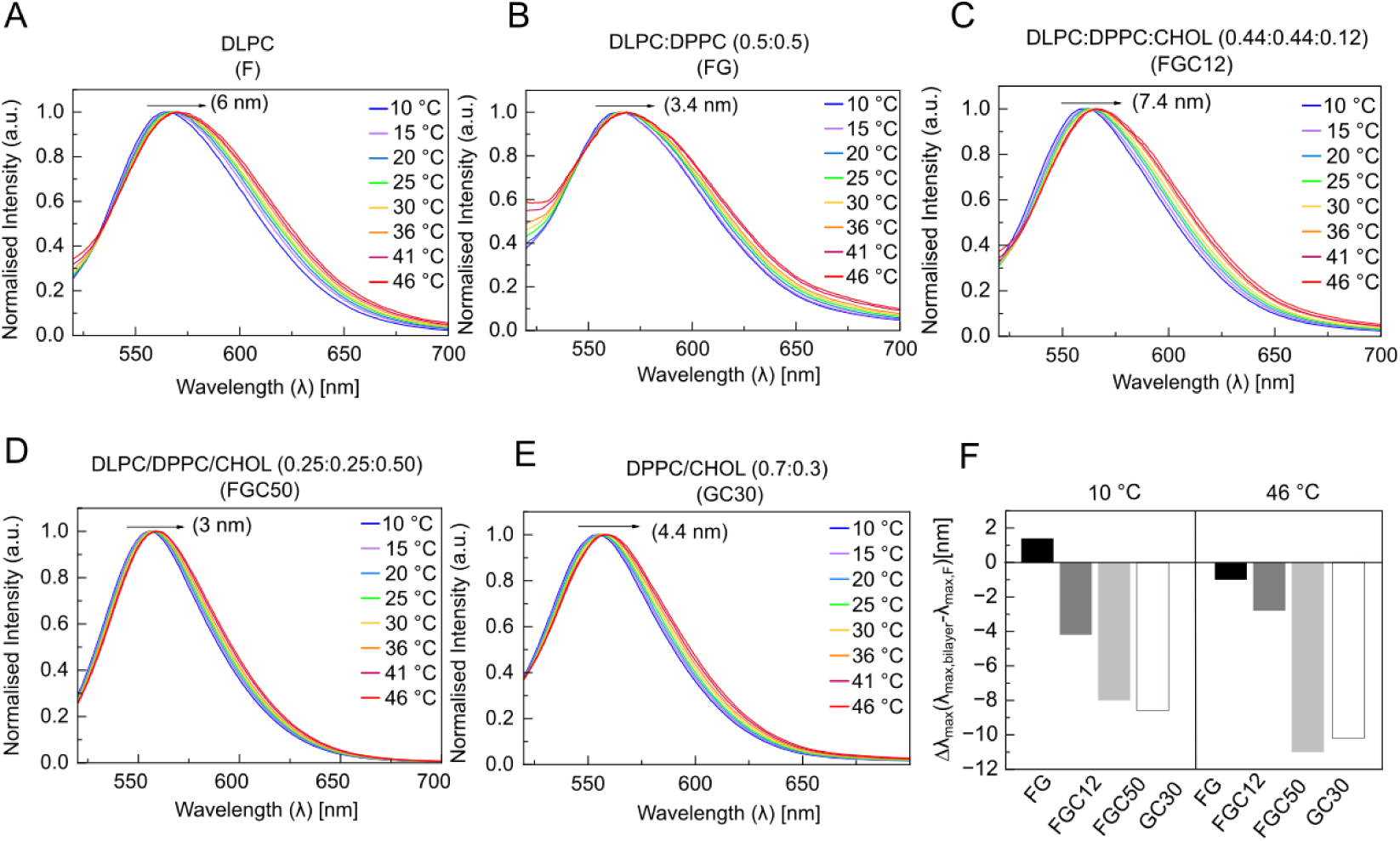
Steady state fluorescence spectra of Di-4 in single, two and three component LUVs within temperature range 10-46°C recorded at λ_ex_ = 476 nm. A) DLPC (F), B) DLPC/DPPC (FG), C) DLPC/DPPC/12 mol% cholesterol (FGC12), D) DLPC/DPPC/50 mol% Cholesterol (FGC50) and E) DPPC/Cholesterol (GC30). F) The difference between 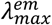 of various bilayers from that of Ld phase F-type bilayer (Δ*λ_max_*) at two extreme experimental temperatures (10°C and 46°C) are shown.

Figs. 2A-E shows the temperature dependence of normalized emission spectra of various Di-4 labelled vesicles. We conducted measurements in 10-46°C range. All vesicles showed slight red shift of the emission maxima 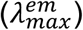 with increasing temperature. While the red shift is not as prominent as that of Laurdan^45,60,61^, another well-established polarity-sensitive probe, in similar context, it indicates general disordering of vesicles at elevated temperature. From the same data set, we evaluated cholesterol dependence of 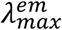 shift since cholesterol is known to induce membrane order. As expected in more ordered and less polar environment, a clear blue shift of 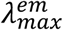 was observed for cholesterol containing *FGC12*, *FGC50*, and *GC30* bilayers compared to *F* type bilayer at a given temperature. A clearer picture of net changes of emission due to temperature or cholesterol content was observed when the 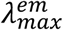 of various bilayers were compared with respect to the *F* bilayer which is the most disordered, most polar and least viscous system tested here. We will do similar comparisons of other parameters throughout the manuscript. The difference of 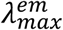 in bilayers compared to *F* type bilayers at the lowest and highest temperatures (10°C and 46°C) are plotted in Fig. 2F. The difference is slightly more prominent at 46°C. However, the difference between 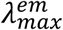 of *F*-type system (most disordered) and *GC30* or *FGC50* type systems (most ordered) is marginal as opposed to those reported earlier (for DOPC/DPPC/Chol systems)^58^. This is likely due to the differences in the lipid type and composition used in these studies. Such small difference in the 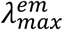 between ordered and disordered states were previously reported in cell derived giant plasma membrane vesicles^45,59^ where the phases are generally less distinctive than DOPC/DPPC/Chol vesicles^27^. Therefore, the small changes in 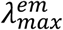 between cholesterol containing bilayers and F type bilayer may be attributed to relatively weak phase separation in the former. Another observation is that 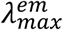 of *F* and *FG* type systems are more or less same; meaning Di-4 does not partition into the gel phase^24^. Overall, we observe modest temperature dependence and slightly stronger cholesterol dependence on the emission spectra of Di-4 in the ternary DLPC/DPPC/Cholesterol system.

### Fluorescence lifetime histogram of Di-4 reveals two regions of differential nanoscale polarity

We next performed the time-resolved measurements on the same bilayers across the same temperature range. We excited the samples at 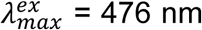 and the time-resolved emission was collected at two emission wavelengths (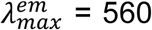 nm and 620 nm). We chose these two emission wavelengths since they are often used to correspond Lo and Ld phases for GP measurements^15,27,62^. However, in our hands, we generally get good signal for TCSPC data analysis at 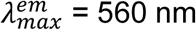. The TCSPC raw data and fits for 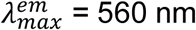 and 620 nm are shown in Figs. 3A-D and S2. The fitted results are given in Table S1 and Table S2. We generally observe similar trends across all bilayers in both the temperature- and cholesterol- dependence measurements for both emission wavelengths (Table S1 and Table S2 and Fig. S4). Therefore, we discuss the results with 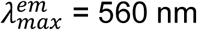 data.

**Figure 3.**
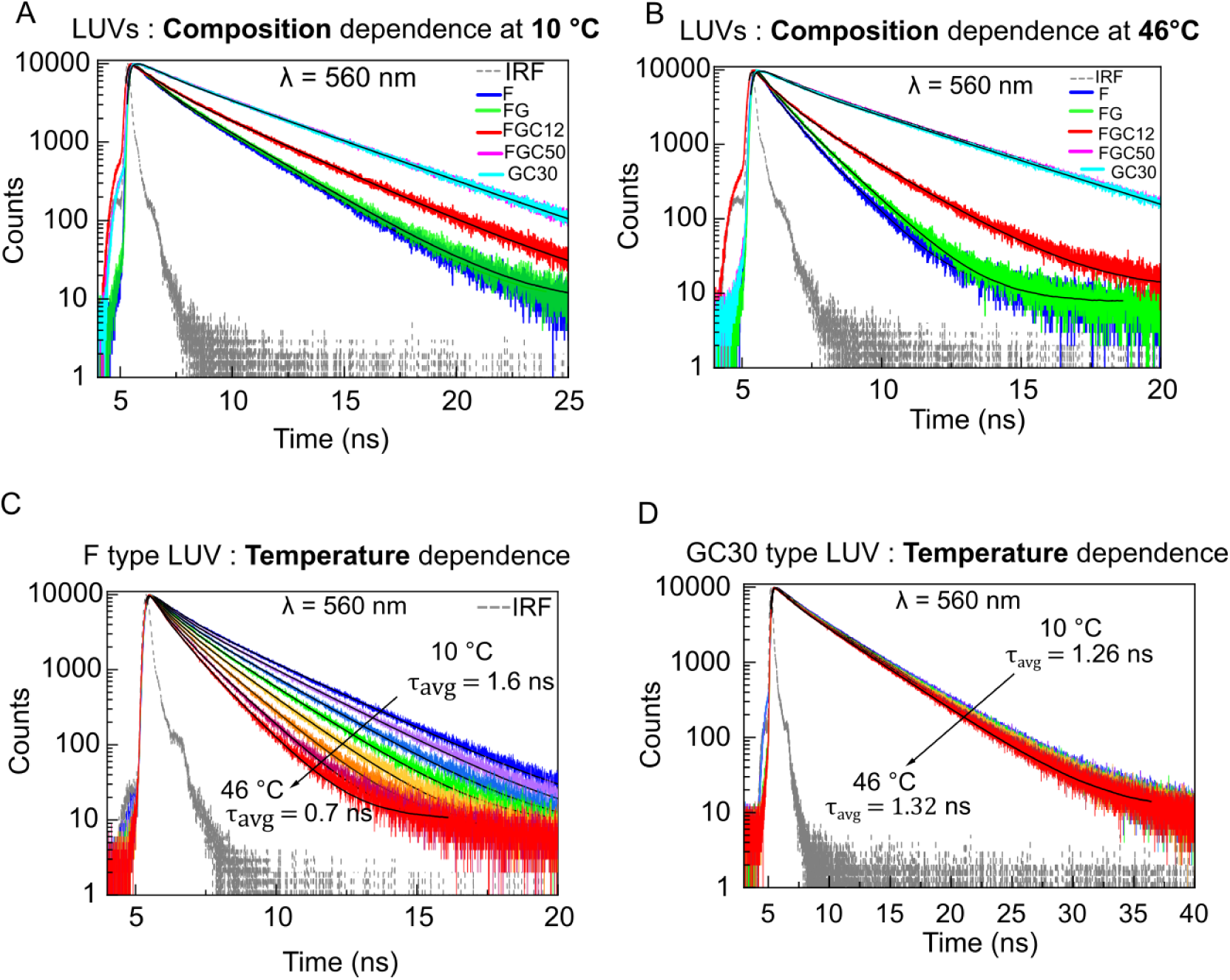
Temperature and composition dependence of Di-4’s fluorescence lifetime. A-B) Raw and fitted fluorescence lifetime histograms of Di-4 on LUVs of various composition (given in Table 1) at two specific temperatures (10° and 46°C). C-D) Raw and fitted fluorescence lifetime histograms of Di-4 on LUVs of F and GC30 LUVs (composition given in Table 1) between 10°C – 46°C. The excitation and emission maxima of raw data recording were 476 nm and 560 nm.

The lifetime histograms of all bilayers (Figs. 3A-D) can be adequately fitted with two-component model (Eq. 1) yielding the long and short lifetime (*τ_long_* and *τ_short_* respectively) values along with their corresponding amplitudes. The average lifetime (*τ_av_*) can be determined from these values using Eq. 2. The *τ_long_* and *τ_short_* values represent two distinctive nanoenviroment around Di-4 while the *τ_av_* mirror the global picture of membrane physical properties sensed by this probe. We observed two- components in the lifetime histogram even for a single component *F*-type bilayer (Table S1). Since the phase transition temperature of DLPC is -2°C,^52,63^ the *F* type bilayer remains in the Ld phase at our experimental conditions. Therefore, distinctive long and short lifetimes of Di-4 may not be attributed to two different phases. A recent study^49^ suggests that the sensor moiety of Di-4 is localized at the surface and hence it described the hydration or polarity level close to the bilayer-water interface. The hydration and polarity are smaller for more tightly packed lipids and vice versa. We therefore infer that Di-4 resides in two different solvation environments (less polar (LP) and more polar (MP)) linked to differential packing. As discussed earlier, we assign the longer lifetime component to less polar (*τ_long_* = *τ_LP_*), and thus more ordered nanoenvironment while the shorter lifetime component arises from more polar (*τ_short_* = *τ_MP_*), and thus more disordered nanoenvironment. Note that the photophysical dynamics given by fluorescence lifetime is in nanosecond timescale (Table S1 and S2) while lateral diffusion of Di-4 in lipid bilayers occur in millisecond timescale^64^. Therefore, the Di-4 molecules remain more or less in the same location during the photophysical processes. Hence, the fluorescence lifetime of this probe reflects the nanoscale polarity of its proximal solvation regions.

### The nanoscale polarity of both more polar (MP) and less polar (LP) regions is influenced by cholesterol

The absolute values of these two lifetime components were different across bilayers (Table S1). Generally, the trend of both *τ_LP_* and *τ_MP_* values across bilayers is: *FGC50* > *GC30* > *FGC12* > *FG* ∼ *F*. Therefore, all bilayers have both LP and MP regions but the degree of polarity of a given region (MP or LP) varies greatly. The melting temperature of DLPC and DPPC lipids is -2 °C and 41 °C respectively^52,63^. It is likely that LP regions, which are more hydrophobic, is enriched with tightly packed DPPC lipids while the MP regions are enriched with relatively less tightly packed DLPC lipids in ternary *FGC* mixtures. Cholesterol is known to impart ‘condensing’ effects on the disordered bilayers and thus induces short-range order. By contrast, it ‘fluidizes’ gel bilayers by disrupting long-range order. Therefore, the extent of polarity in the MP and LP regions in the *FGC* bilayer would depend on cholesterol concentration in these regions. In the same line, the LP regions in *GC* bilayer would have relatively less concentration of cholesterol than in the MP region given the ‘fluidizing effects’ of cholesterol on a gel supporting lipid bilayer.

Unlike steady-state spectra (Fig. 2), lifetime histogram of Di-4 for all bilayers is strongly temperature sensitive (Fig. 3). The contrast between steady-state and time-resolved spectra reflects temperature dependent ultrafast dynamics of the nano- environment sensed by Di-4. We show the raw lifetime histograms and their corresponding fits of different bilayers only at two extreme experimental temperatures (10°C and 46 °C) in Figs. 3A and 3B while the fitted data at other temperatures are given in Table S1. The average fluorescence lifetime (*τ_av_*) of Di-4 in F type bilayer decreases from 1.6 ns at 10°C to 0.7 ns at 46°C (Fig. 3C). These lifetime values are comparable to the literature reports on similar systems^65^. This bilayer remains in the fluid phase in the experimental temperature range (10-46°C). Therefore, the change in the lifetime is not due to changes in phase state. The thermal energy in the system at higher temperatures makes the lipid acyl chains more dynamic and hence overall membrane becomes more disordered and hence more polar. This facilitates excited state torsional rotation of the Di-4 which increases the contribution of non-radiative process in the overall excited state relaxation of the molecules leading to a decrease in the *τ_av_* value. This interpretation corresponds well with the diffusion measurements on DLPC bilayer^66^. The disordering of the acyl chain at higher temperature also increased the lateral diffusion of the lipid probes.

The temperature dependence of cholesterol containing bilayers were rather weaker as observed from the fluorescence lifetime histograms of F and GC type bilayers (Fig. 3C and D and Table S1). The trends in the changes of *τ_av_* value by cholesterol content may also be explained in a similar fashion as above. Since cholesterol is known to order the plasma membrane, a larger concentration of this component leads to restriction of the aforementioned torsional rotation and thereby increasing *τ_av_* (Fig. S3 and Table S1).

We next examined the temperature dependence of *τ_LP_* and *τ_MP_* for different bilayers. Both *τ_LP_* and *τ_MP_* values, similar to *τ_av_* values, decrease with temperature (Table S1) for the same reason stated earlier for the *τ_av_* trend. To gain better insights into the molecular organization across various bilayers, we determine the ratio of *τ_LP_* of a given bilayer to that of *F*-type bilayer. This ratio, 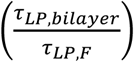, is the spectroscopic descriptor of nanoscale polarity of LP membrane region of a given bilayer relative to the LP region of liquid disordered bilayer. The analogous descriptor for MP membrane environment is the ratio 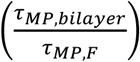. We name these ratios ‘relative index of nanoscale polarity’ (*RINP*) of LP and MP regions (*RINP_LP_* and *RINP_MP_* respectively) and their values are 1 for *F*-type bilayer. The *RINP* values are inversely proportional to the nanoscale polarity. For the *FG* bilayer, both *RINP_LP_* and *RINP_MP_* values are close to 1 (Figs. 4A and 4B) indicating that Di-4 fails to sense the changes in membrane order due to incorporation of saturated lipids. By contrast, both *RINP_LP_* and *RINP_MP_* values are greater than 1 in the cholesterol containing *FGC* and *GC* bilayers. This means that cholesterol makes both LP and MP regions less polar and more hydrophobic. Since *FGC* and *GC* bilayers have Lo-Ld and Lo phases while *FG* bilayer has coexisting So- Ld phase,^67^ it is clear that Di-4 does not partition into the So state (*vide supra*). Di-4 only senses the Ld region of the *FG* bilayer and hence its behaviour resembles to that of a *F* bilayer.

**Figure 4.**
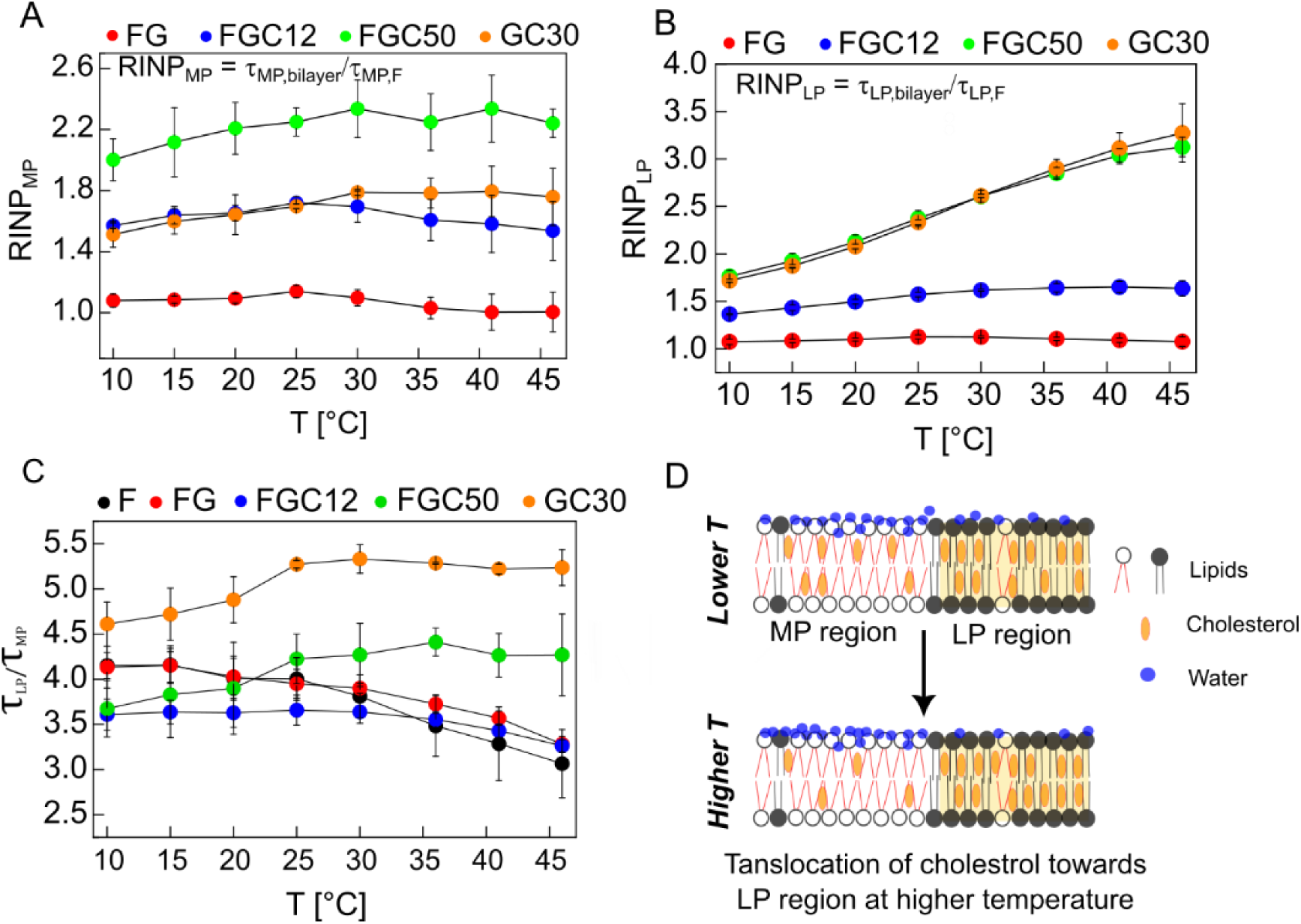
Relative polarity of MP and LP regions and their temperature dependence across various bilayer and within same bilayer as determined from Di-4 lifetime components. A) Temperature variation of ‘Relative Index of Nanoscale Polarity’ of MP region (RINP_MP_), given by the ratio of ratio of *τ_MP_* of a given bilayer (*FG*, *GC30*, *FGC12*, *FGC50*) to that of the *F* bilayer 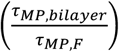, across various temperature. B) Temperature variation of RINP of LP region (RINP_LP_), given by the ratio of ratio of *τ_LP_* of a given bilayer (*FG*, *GC30*, *FGC12*, *FGC50*) to that of the *F* bilayer 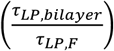, across various temperature. C) The relative polarity of the LP and MP regions within the same bilayer is given by the ratio (*τ_LP_*/*τ_MP_*) which is plotted for various bilayers (*F*, *FG*, *GC30*, *FGC12*, *FGC50*) and its temperature dependence. D) A graphical representation pictorially explaining cholesterol translocation from MP to LP regions at higher temperatures as a possible mechanism of the observed trends in Figs. 4A-C.

We plotted *RINP_MP_* and *RINP_LP_* against temperature for all bilayers (Figs. 4A and 4B). We observed that the values of both the indices increase in the presence of cholesterol. The *RINP_MP_* value increases as: *FGC50* > *FGC12* ∼ *GC30* > *FG* ∼ 1. This indicates that the hydrophobicity of the more polar regions increases upon introduction of cholesterol in the system. The *RINP_MP_* values for various bilayers were more or less temperature independent. This means that temperature dependent changes of cholesterol induced ordering in the MP regions occur similarly across various bilayers. Interestingly, this was not the case for LP regions. Here we see clearly the trends differ strongly between Lo containing *FGC-50* and *GC-30* bilayers and relatively less ordered *FGC-12* bilayer. The *RINP_LP_* values increase with temperature for FGC-50 and GC-30 bilayers. This means that the rate of polarity change with temperature in the LP regions of these two systems is somewhat less prominent than in disordered F system. We reasoned that at higher temperatures the LP regions of FGC-50 and GC-30 systems have higher cholesterol content than lower temperature, making this region even less polar (Fig. 4D). Increase in cholesterol content may be either due to the translocation of cholesterol from the MP region to the LP region on the outer leaflet or cholesterol may also be flipped from the inner leaflet^68^. Note that Di-4 is localized in the outer leaflet only^69^. In any case, the polarity difference between the LP and MP regions of FGC-50 and GC-30 systems is greater at higher temperature. This is supported from the observed larger values of *τ_LP_*/*τ_MP_* at higher temperature for these two systems (Fig. 4C). These interesting cholesterol dependent features (Fig. 4D) was only observed in systems where cholesterol content is high. In the low cholesterol FGC-12 system the temperature dependent movement of cholesterol, if at all happens, would not sufficiently change polarity of LP and MP regions to be observed by our analysis.

### The Excited State Relaxation Activation Energy (ESRAct) of Di-4 can be determined from the temperature dependence of its lifetime

The rate constant of excited state relaxation is inversely proportional to fluorescence lifetime^70^. We determine this rate constant for LP and MP regions (*k_LP_* and *k_MP_* respectively) at each temperature by taking inverse of *τ_LP_* and *τ_MP_* respectively. We found out that the dependence of both *k_LP_*(= 1/*τ_LP_*) and *k_MP_*(= 1/*τ_MP_*) is Arrehenius-like (Fig. 5), and not sigmoidal, within the temperature range (10- 46°C). The reported phase transition temperatures of *FGC* and *FG* type bilayers are within this range^50,67,71,72^. Therefore, temperature dependence of Di-4’s lifetime does not reflect phase transition of these bilayers. This further confirms photophysical properties of Di-4 are only sensitive to local heterogeneity; rather than global phase transition.

**Figure 5.**
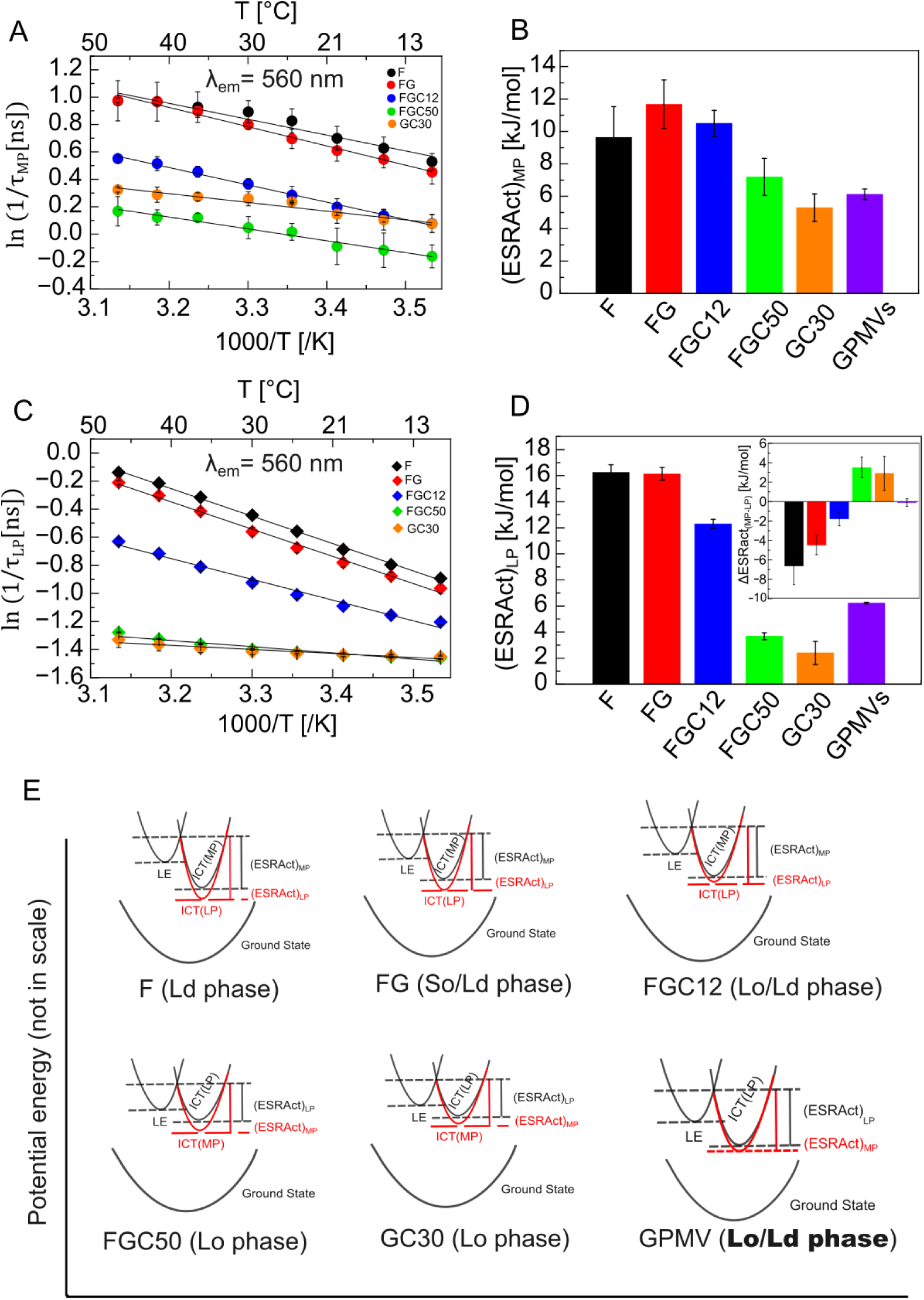
Arrhenius plots of Di-4 and the corresponding ESRAct values of various LUVs and MCF-7 derived GPMVs. (A-B) MP region, and (C-D) LP region. The difference between ESRAct values of MP and LP regions, ΔESRAct_(MP-LP)_ or simply ΔESRAct, for different systems in the inset of Fig. D. E) The model of potential energy diagram of various LUV systems based on the results in Fig. A-D.

The temperature dependence of both 1/*τ_LP_* and 1/*τ_MP_* is fitted with Arrhenius equation (Eq. 3) to determine activation energy of these processes. The values were in the following order: *F* ∼ *FG* > *FGC12* > *FGC50* ∼ *GC30* (Figs. 5A-D). Since Di-4 is sensitive to solvent polarity, it may be thought that the water dynamics around solvated Di-4 is strongly temperature-sensitive. It was reported that the activation energy of water diffusion is in the order of 20 kJ/mol^73^ which is larger than the activation energy of the dynamics reflected by the temperature dependence of Di-4’s fluorescence lifetime (5-16 kJ/mol, Figs. 5A-D). Therefore, we rule out the activation energy obtained here is due to diffusion dynamics of water surrounding Di-4 molecules. Rather, this activation energy appears to correspond to excited state relaxation processes and hence this energy barrier is called excited state relaxation activation energy (ESRAct). The excited state relaxation can occur via both radiative and non-radiative pathways. Generally, the radiative rate constant is temperature independent^70^. Therefore, the observed activation energy is likely for the non-radiative processes. In the current context, a prominent non-radiative process is the transition between LE and ICT states which is known to require an activation energy^74^. The dynamics of this non-radiative transition, as discussed earlier, is influenced by the surrounding solvation layer which varies across different bilayer systems. The presence of water layer, as in the case of *F* and *FG* type bilayers, allowed torsional rotation of the excited Di-4 molecules leading to the stabilization of the ICT state. When cholesterol is added to this system it occupies the interstitial spaces (evident from the increase of both *τ_LP_* and *τ_MP_* upon cholesterol addition; Table S1 and Table S2) replacing the water molecules leading to the reduction of overall hydration and polarity. Therefore, the ICT states in cholesterol-containing systems (ICT_FGC50_ and ICT_GC30_) have energy relatively closer to the LE state. Overall, ICT states have lower energy than the LE state in general for all bilayers. However, the ICT states of *FGC50* and *GC30* type bilayers have higher energy than those *F* and *FG* of type bilayer. So, the LE state is closer to the energy levels of both ICT_FGC50_ and ICT_GC30_ states while ICT_F_ and ICT_FG_ states have much lower energy.

### ESRAact of Di-4 of MP and LP regions of various bilayers corroborates with their respective phase separation

We observed that the ESRAct values for both LP and MP regions of the cholesterol-containing systems (*FGC50* and *GC30*) are much smaller than the *F* and *FG* systems (Figs. 5B and 5D). This means that the activation energy of LE ↔ ICT_FGC50_ and LE ↔ ICT_GC30_ transition is smaller than LE ↔ ICT_F_ and LE ↔ ICT_FG_ transition (Fig. 5E). Therefore, while ICT_F_ and ICT_FG_ states are energetically more stable than the LE states, the Di-4 molecules are required to overcome a larger energy barrier. This means that, on average, there is a relatively higher probability of emission from the LE state of *F*, *FG*, and *FGC12* systems than the *FGC50* and *GC30* systems^75^. This is also probably the reason behind the observed small difference in the *λ^em^* between *F* and *FGC50* or *GC30* bilayers. The ESRAct of Di-4 therefore clearly differentiates cholesterol dependent physical states of lipid vesicles; which complements the phase diagram of the ternary lipid mixtures^50^.

A closer look at the ESRAct values for LP and MP regions across different bilayers reveals an interesting feature. The ESRAct of F and FG bilayers at LP regions is larger than in the MP regions. By contrast, the same of the *FGC50* and *GC30* bilayer at LP region is smaller than in the MP regions. The ΔESRAct between MP and LP regions are negative for *F*, *FG*, and *FGC12* bilayers while it is positive for *FGC50* and *GC30* bilayers (Fig. 5D, inset). This may be rationalized as follows (Fig. 5E): In *F*, *FG,* and *FGC12* bilayers, MP regions are less tightly packed and thus have more packing defects into which the water molecules can penetrate. By contrast, the LP regions have more tightly packed lipids making this ICT_LP_ states more energetically stable and thus ICT_LP_ ↔ LE transition has larger activation energy. The scenario is different in the *FGC50* and *GC30* bilayers. Here, the lipid packing is disturbed by the cholesterol. In addition, the MP region has more water molecules while LP region has more cholesterol molecules. This means that Di-4’s torsional rotation is more facilitated in the MP regions, making ICT_MP_ energetically more stable than the ICT_LP_ state and hence larger activation energy is required for ICT_MP_ ↔ LE transition. The radical difference between the underlying molecular packing and organization is reflected in the net differences of ESRAct values between MP and LP regions.

### ESRAct of Di-4 reveals Lo-Ld co-existing phase separation on MCF-7 derived giant plasma membrane vesicles (GPMVs)

We applied the principles of ESRAct on MCF-7 derived giant plasma membrane vesicles (GPMVs) of average size ∼15 μm (Fig. 6A). GPMVs more or less maintain the composition of the plasma membrane they are derived from. The confocal image of Di-4 labelled GPMVs does not show any obvious phase separation at room temperature (22°C). The fluorescence lifetime of Di-4 in GPMVs however shows moderate temperature dependence within the range 4–37°C (Fig. 6B and Table S3).

**Figure 6.**
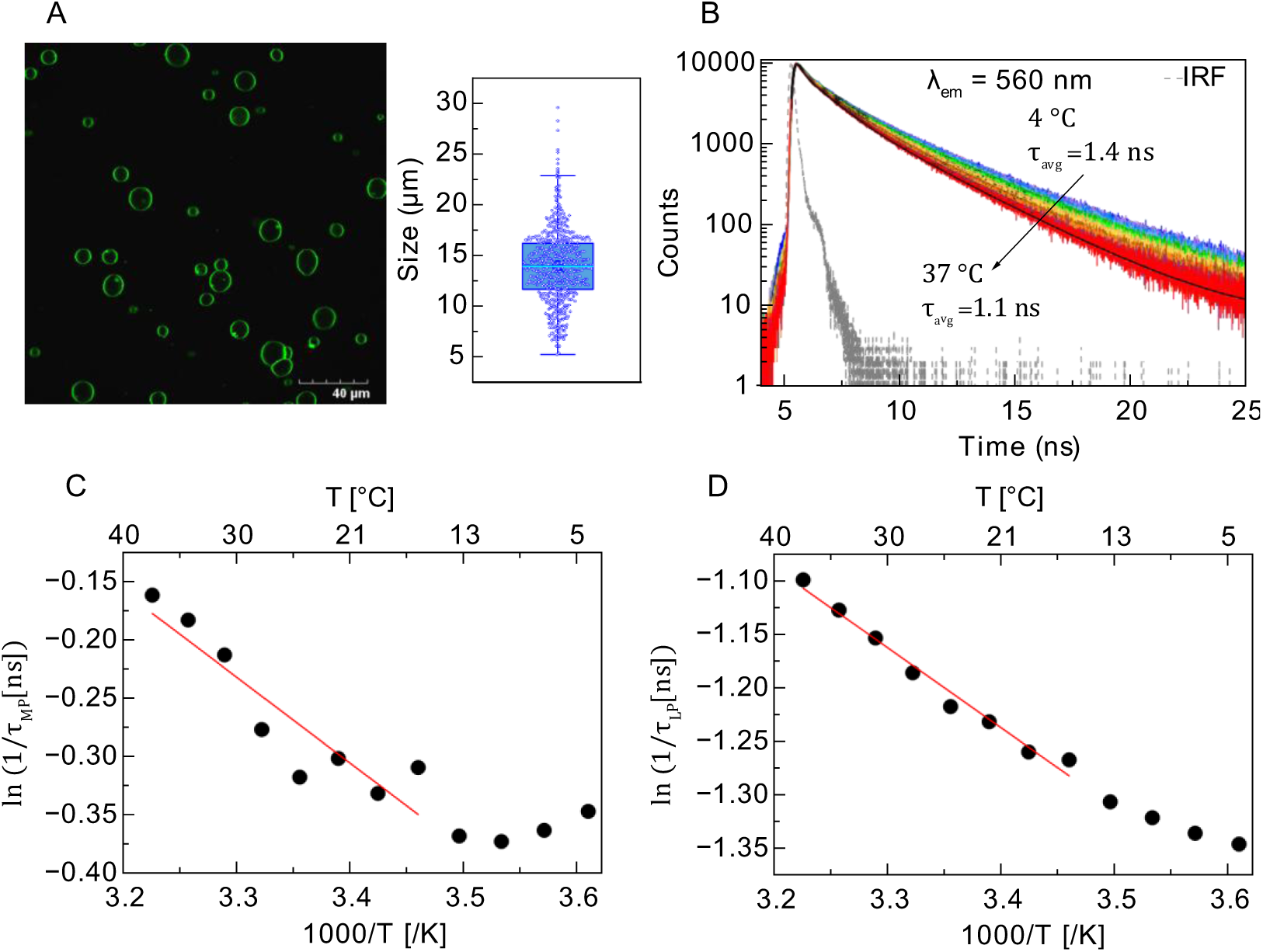
ESRAct analysis of MCF-7 derived GPMVs. A) Confocal images of Di-4 labelled GPMVs and corresponding size distributions. B) Fluorescence lifetime histograms and their fits of Di-4 labelled GPMVs are taken between temperature range 10° to 37 °C. C, D) Arrhenius plots associated to the excited state relaxation at MP and LP regions. The excitation and emission maxima of raw data recording were 476 nm and 560 nm.

The temperature dependence of Di-4 lifetime in GPMVs is contrasting to the Ld phase bilayer where we observed strong temperature dependence of Di-4 lifetime. The Arrhenius plot for the MP region clearly show two distinct of temperature dependence (Fig. 6C): in the 4–13°C range, the lifetime of Di-4 is almost temperature independent while modest temperature dependence was observed in the 16–37°C range. This feature in not observed on the LUVs (Fig. 5) or the LP regions of GPMVs (Fig. 6D). It is known that the phase mixing temperature (*T*_misc_) of GPMVs prepared using the PFA/DTT method used here occurs within 15-20°C^76–79^. At temperatures higher than *T*_misc_ micron-sized ordered regions (microdomains) transform into nanoscale ordered regions (nanodomains) which is present at physiological temperature^34^. Our observations in the MP regions might reflect that although other possibilities such as differences in the protein solvation may not be ruled out. Nonetheless, we determined the ESRAct values for LP and MP regions from the data at 16–37°C range. The ESRAct values in both regions were very close (∼6.2 kJ/mol) and thus ΔESRAct between MP and LP regions is around zero. When compared to the LUV systems, these values fall in between *FGC12* and *FGC50*/*FG30* systems. We therefore conclude that MCF-7 derived GPMVs exhibit Lo-Ld phase separation in the 16–37°C range.

## CONCLUSION

In this study, we used ESRAct of Di-4 from the temperature dependence of its fluorescence lifetime to understand membrane phase state. We found out that there are both more polar and less polar regions in all bilayers regardless of their phase state. However, the ESRAct values of Di-4 in both regions are correlated with their phase states. Generally, the ESRAct values decreases going from Ld ◊ Lo/Ld ◊ Lo phases in the lipid bilayer. The ESRAct value of GPMVs derived from MCF-7 cell lines resembles that of Lo/Ld state. Such information was not possible to gain from steady-state measurements of this dye.

Overall, this study systematically establishes the principles of ESRAct of a polarity sensitive dye using various compositions of a ternary mixture and ranks these values with respect to the nanoscale phase separation. ESRAct can be used as photophysical and nearly non-invasive metric to evaluate physical states of biological systems at various experimental conditions. This straightforward method requires only temperature dependence lifetime measurements. These principles may easily be extended to fluorescence lifetime imaging (FLIM) modality as well as to other polarity sensitive probes. We envisage wide applications of this method in biophysical and photophysical investigations in future.

## ASSOCIATED CONTENTS

### Supporting Information

Four figures and three tables including the analyzed data of time-resolved fluorescence measurements (PDF).

## AUTHOR INFORMATION

### Notes

The authors declare no competing interest.

## Author Contributions

D.M. and N.B. conceived the project and designed the experiments; D.M. and A.T. preformed the experiments; D.M and N.B. analyzed data and wrote the manuscript with comments from A.T.

## ACKNOWLEDGEMENTS

D.M. acknowledges the PhD research fellowship from IIT Kharagpur. A.T. acknowledges DST-INSPIRE research fellowship and support. All authors thank Prof. Nilmoni Sarkar (IIT Kharagpur) for the cell culture facility. The research facilities of the Department of Chemistry, IIT Kharagpur are greatly acknowledged. N.B. acknowledges the funding from IIT Kharagpur (IIT/SRIC/CY/IIT/2023–2024/162) and SRG fund from Anusandhan National Research Foundation (ANRF) (SRG/2023/001423).

