## Supplemental Information for "Excited State Relaxation Activation Energy (ESRAct) of Di-4-ANEPPDHQ Maps Nanoscale Molecular Organization in Biomembranes"

**FIGURE S1**

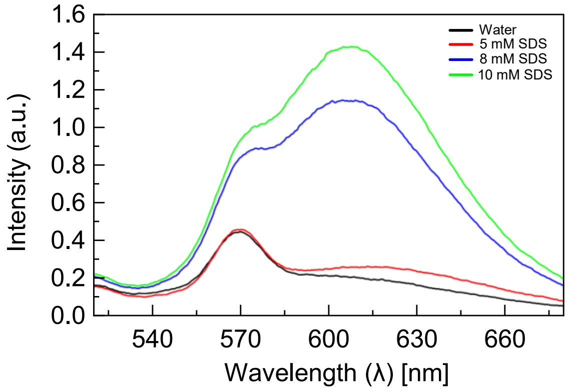


**Figure S1.** Emission spectra of Di-4 recorded at $\lambda_{ex}$ = 476 nm in water and different concentrations of SDS surfactants at [SDS] < $[SDS]_{CMC}$ and [SDS] > $[SDS]_{CMC}$. $[SDS]_{CMC}$ = 8 mM[1].

**FIGURE S2**

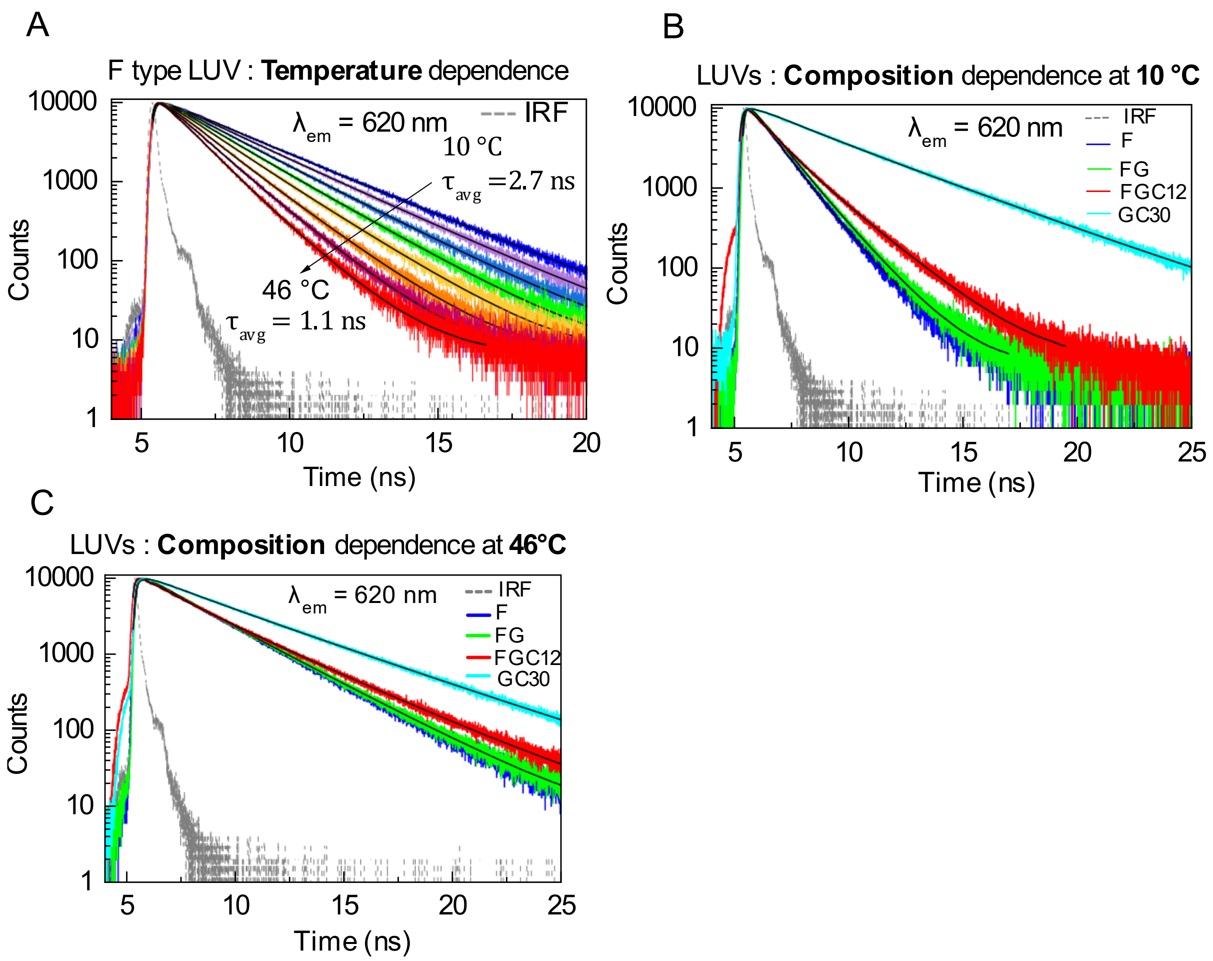


**Figure S2:** Variation of Di-4 lifetime as a function of temperature and lipid composition. A) Raw and fitted fluorescence lifetime histograms of Di-4 in F type LUVs in the 10° to 46 °C temperature range. B-C) Variation of fluorescence lifetime histograms among LUVs of different composition at 10°C (B) and 46°C (C). The excitation and emission maxima of raw data recording were 476 nm and 620 nm.

**FIGURE S3**

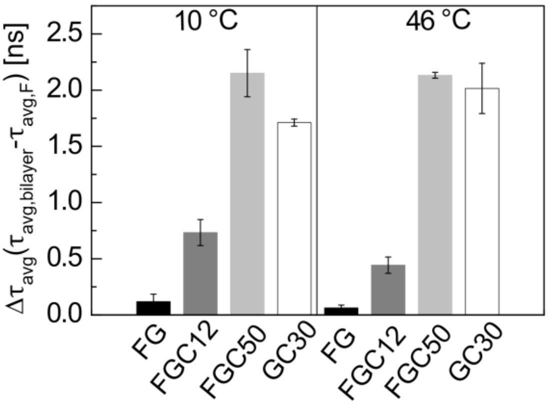


**Figure S3:** Difference of average fluorescence lifetime at $\lambda_{em}$ = 560 nm of different LUVs in comparison to *F* type LUVs at 10°C and 46°C.

**FIGURE S4**

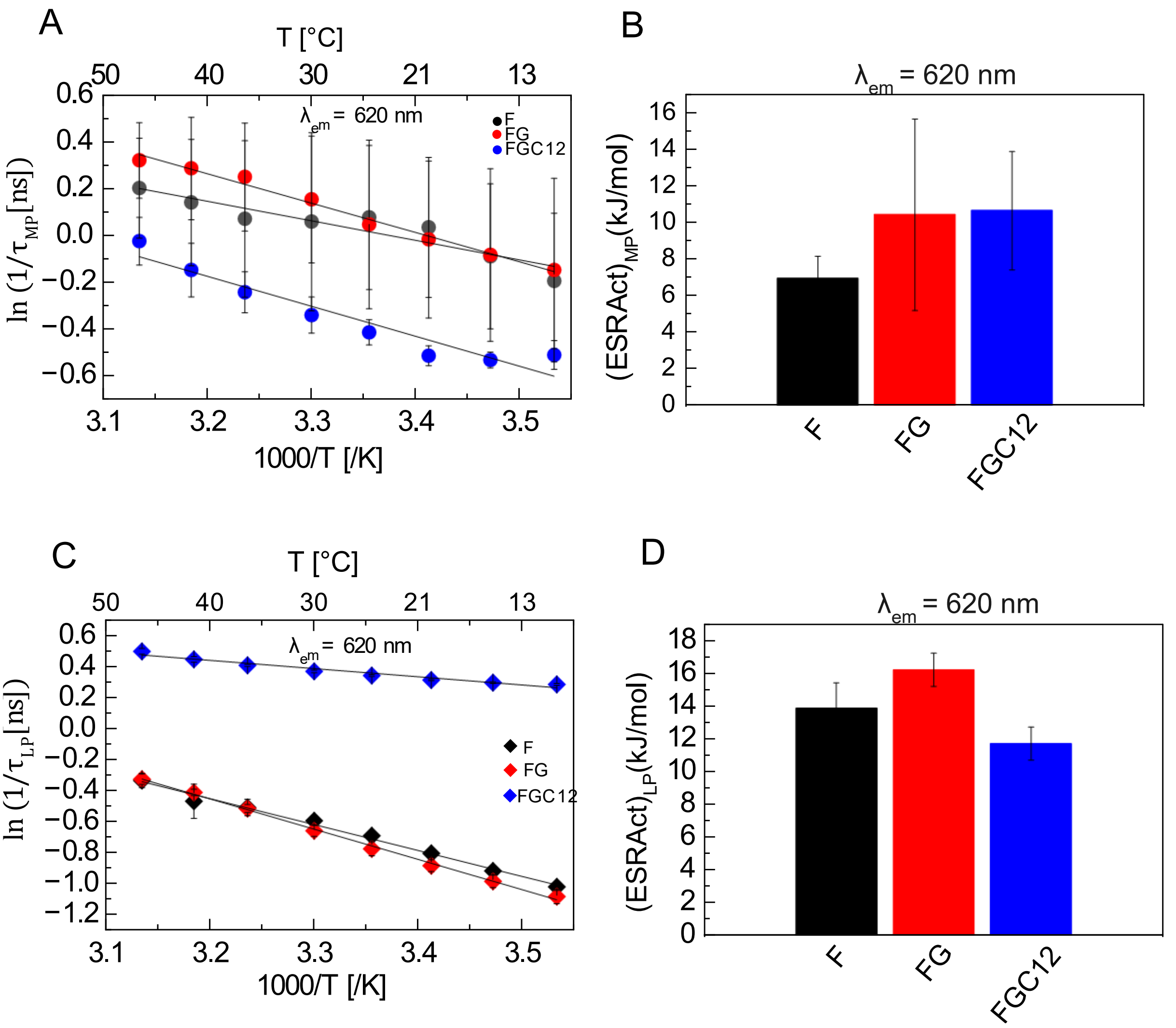


**Figure S4.** Arrhenius plots and ESRAct values of Di-4 in the more polar (MP) and less polar (LP) regions of different LUVs recorded at $\lambda_{em}$ = 476 nm and $\lambda_{em}$ = 620 nm. A-B) MP region, and C-D) LP region of *F, FG, and FGC12* type lipid vesicles.

Table S1: Values of the fit parameters of fluorescence lifetime histograms across different temperature and composition of LUVs at $\lambda_{em}$ = 560 nm.

| LUV | Temp. [ºC] | $\tau_{MP}\ [ns]$ | $\alpha_{MP}$ | $\tau_{LP}\ [ns]$ | $\alpha_{LP}$ | $\tau_{Avg}\ [ns]$ |
|---|---|---|---|---|---|---|
| F | 10 | 0.59 ± 0.03 | 0.49 ± 0.06 | 2.44 ± 0.06 | 0.51 ± 0.06 | 1.54 ± 0.16 |
| | 15 | 0.53 ± 0.04 | 0.51 ± 0.06 | 2.22 ± 0.07 | 0.49 ± 0.06 | 1.36 ± 0.16 |
| | 20 | 0.49 ± 0.04 | 0.53 ± 0.06 | 1.99 ± 0.05 | 0.47 ± 0.06 | 1.20 ± 0.14 |
| | 25 | 0.44 ± 0.04 | 0.54 ± 0.06 | 1.75 ± 0.05 | 0.46 ± 0.06 | 1.05 ± 0.12 |
| | 30 | 0.41 ± 0.03 | 0.55 ± 0.06 | 1.56 ± 0.03 | 0.45 ± 0.06 | 0.93 ± 0.09 |
| | 36 | 0.39 ± 0.04 | 0.57 ± 0.05 | 1.37 ± 0.02 | 0.43 ± 0.05 | 0.81 ± 0.08 |
| | 41 | 0.38 ± 0.05 | 0.59 ± 0.03 | 1.24 ± 0.02 | 0.41 ± 0.03 | 0.73 ± 0.07 |
| | 46 | 0.38 ± 0.05 | 0.64 ± 0.03 | 1.15 ± 0.04 | 0.36 ± 0.03 | 0.66 ± 0.05 |
| | Temp. [ºC] | $\tau_{MP}\ [ns]$ | $\alpha_{MP}$ | $\tau_{LP}\ [ns]$ | $\alpha_{LP}$ | $\tau_{Avg}\ [ns]$ |
| FG | 10 | 0.64 ± 0.05 | 0.48 ± 0.04 | 2.62 ± 0.09 | 0.51 ± 0.04 | 1.66 ± 0.14 |
| | 15 | 0.58 ± 0.03 | 0.49 ± 0.04 | 2.40 ± 0.06 | 0.51 ± 0.04 | 1.51 ± 0.12 |
| | 20 | 0.54 ± 0.04 | 0.50 ± 0.04 | 2.18 ± 0.07 | 0.49 ± 0.04 | 1.35 ± 0.12 |
| | 25 | 0.49 ± 0.03 | 0.51 ± 0.05 | 1.97 ± 0.06 | 0.48 ± 0.05 | 1.21 ± 0.12 |
| | 30 | 0.45 ± 0.01 | 0.53 ± 0.06 | 1.75 ± 0.05 | 0.46 ± 0.06 | 1.06 ± 0.11 |
| | 36 | 0.41 ± 0.02 | 0.55 ± 0.07 | 1.51 ± 0.04 | 0.44 ± 0.07 | 0.90 ± 0.09 |
| | 41 | 0.38 ± 0.01 | 0.57 ± 0.07 | 1.35 ± 0.02 | 0.42 ± 0.07 | 0.79 ± 0.08 |
| | 46 | 0.37 ± 0.006 | 0.59 ± 0.07 | 1.23 ± 0.01 | 0.40 ± 0.07 | 0.72 ± 0.06 |
| | Temp. [ºC] | $\tau_{MP}\ [ns]$ | $\alpha_{MP}$ | $\tau_{LP}\ [ns]$ | $\alpha_{LP}$ | $\tau_{Avg}\ [ns]$ |
| FGC12 | 10 | 0.93 ± 0.06 | 0.44 ± 0.01 | 3.33 ± 0.07 | 0.55 ± 0.01 | 2.27 ± 0.07 |
| | 15 | 0.87 ± 0.04 | 0.45 ± 0.01 | 3.18 ± 0.05 | 0.54 ± 0.01 | 2.11 ± 0.05 |
| | 20 | 0.82 ± 0.05 | 0.48 ± 0.01 | 2.98 ± 0.07 | 0.52 ± 0.01 | 1.94 ± 0.04 |
| | 25 | 0.75 ± 0.05 | 0.49 ± 0.02 | 2.75 ± 0.07 | 0.50 ± 0.02 | 1.75 ± 0.04 |
| | 30 | 0.69 ± 0.02 | 0.51 ± 0.02 | 2.52 ± 0.04 | 0.48 ± 0.02 | 1.58 ± 0.03 |
| | 36 | 0.63 ± 0.02 | 0.54 ± 0.02 | 2.25 ± 0.06 | 0.46 ± 0.02 | 1.38 ± 0.03 |
| | 41 | 0.59 ± 0.03 | 0.56 ± 0.02 | 2.05 ± 0.07 | 0.43 ± 0.02 | 1.23 ± 0.04 |
| | 46 | 0.57 ± 0.01 | 0.59 ± 0.02 | 1.88 ± 0.04 | 0.40 ± 0.02 | 1.10 ± 0.03 |
| | Temp. [ºC] | $\tau_{MP}\ [ns]$ | $\alpha_{MP}$ | $\tau_{LP}\ [ns]$ | $\alpha_{LP}$ | $\tau_{Avg}\ [ns]$ |
| FGC50 | 10 | 1.18 ± 0.10 | 0.19 ± 0.01 | 4.30 ± 0.05 | 0.80 ± 0.01 | 3.69 ± 0.07 |
| | 15 | 1.13 ± 0.14 | 0.20 ± 0.01 | 4.27 ± 0.04 | 0.79 ± 0.01 | 3.62 ± 0.06 |
| | 20 | 1.10 ± 0.15 | 0.21 ± 0.01 | 4.23 ± 0.03 | 0.78 ± 0.01 | 3.54 ± 0.06 |
| | 25 | 0.98 ± 0.06 | 0.22 ± 0.01 | 4.14 ± 0.02 | 0.77 ± 0.01 | 3.43 ± 0.03 |
| | 30 | 0.96 ± 0.08 | 0.24 ± 0.01 | 4.06 ± 0.01 | 0.76 ± 0.01 | 3.31 ± 0.05 |
| | 36 | 0.89 ± 0.02 | 0.25 ± 0.01 | 3.91 ± 0.01 | 0.74 ± 0.01 | 3.15 ± 0.03 |
| | 41 | 0.88 ± 0.05 | 0.26 ± 0.01 | 3.77 ± 0.005 | 0.73 ± 0.01 | 3.00 ± 0.03 |
| | 46 | 0.85 ± 0.09 | 0.29 ± 0.01 | 3.59 ± 0.01 | 0.70 ± 0.01 | 2.79 ± 0.07 |
| | Temp. [ºC] | $\tau_{MP}\ [ns]$ | $\alpha_{MP}$ | $\tau_{LP}\ [ns]$ | $\alpha_{LP}$ | $\tau_{Avg}\ [ns]$ |
| GC30 | 10 | 0.93 ± 0.06 | 0.27 ± 0.01 | 4.27 ± 0.05 | 0.73 ± 0.01 | 3.37 ± 0.02 |
| | 15 | 0.90 ± 0.06 | 0.28 ± 0.01 | 4.25 ± 0.05 | 0.72 ± 0.01 | 3.31 ± 0.03 |
| | 20 | 0.86 ± 0.06 | 0.28 ± 0.005 | 4.21 ± 0.06 | 0.71 ± 0.005 | 3.26 ± 0.04 |
| | 25 | 0.79 ± 0.004 | 0.29 ± 0.005 | 4.16 ± 0.05 | 0.70 ± 0.005 | 3.15 ± 0.02 |
| | 30 | 0.77 ± 0.04 | 0.31 | 4.12 ± 0.09 | 0.69 | 3.07 ± 0.08 |
| | 36 | 0.76 ± 0.02 | 0.32 ± 0.005 | 4.02 ± 0.12 | 0.67 ± 0.005 | 2.97 ± 0.12 |
| | 41 | 0.75 ± 0.04 | 0.33 ± 0.01 | 3.92 ± 0.17 | 0.66 ± 0.01 | 2.85 ± 0.18 |
| | 46 | 0.72 ± 0.01 | 0.35 ± 0.02 | 3.79 ± 0.20 | 0.65 ± 0.02 | 2.71 ± 0.21 |

Table S2: Values of the fit parameters of fluorescence lifetime histograms across different temperature and lipid composition of LUVs at $\lambda_{em}$ = 620 nm.

| LUV | Temp. [ºC] | $\tau_{MP}$ $[ns]$ | $\alpha_{MP}$ | $\tau_{LP}$ $[ns]$ | $\alpha_{LP}$ | $\tau_{Avg}$ $[ns]$ |
|---|---|---|---|---|---|---|
| F | 10 | 1.26 ± 0.32 | 0.16 ± 0.08 | 2.77 ± 0.06 | 0.84 ± 0.08 | 2.50 ± 0.23 |
| | 15 | 1.14 ± 0.31 | 0.17 ± 0.08 | 2.50 ± 0.06 | 0.83 ± 0.08 | 2.25 ± 0.22 |
| | 20 | 1.00 ± 0.26 | 0.17 ± 0.08 | 2.23 ± 0.05 | 0.82 ± 0.08 | 2.00 ± 0.19 |
| | 25 | 0.96 ± 0.26 | 0.19 ± 0.07 | 1.99 ± 0.04 | 0.80 ± 0.07 | 1.78 ± 0.16 |
| | 30 | 1.00 ± 0.32 | 0.29 ± 0.01 | 1.81 ± 0.05 | 0.70 ± 0.01 | 1.57 ± 0.13 |
| | 36 | 0.98 ± 0.29 | 0.47 ± 0.15 | 1.66 ± 0.09 | 0.52 ± 0.15 | 1.36 ± 0.09 |
| | 41 | 0.89 ± 0.21 | 0.58 ± 0.18 | 1.60 ± 0.18 | 0.42 ± 0.18 | 1.19 ± 0.06 |
| | 46 | 0.83 ± 0.16 | 0.61 ± 0.14 | 1.39 ± 0.06 | 0.38 ± 0.14 | 1.07 ± 0.04 |
| | Temp. [ºC] | $\tau_{MP}$ $[ns]$ | $\alpha_{MP}$ | $\tau_{LP}$ $[ns]$ | $\alpha_{LP}$ | $\tau_{Avg}$ $[ns]$ |
| FG | 10 | 1.24 ± 0.41 | 0.20 ± 0.01 | 2.95 ± 0.13 | 0.79 ± 0.01 | 2.59 ± 0.21 |
| | 15 | 1.15 ± 0.36 | 0.18 ± 0.02 | 2.68 ± 0.11 | 0.81 ± 0.02 | 2.39 ± 0.19 |
| | 20 | 1.07 ± 0.31 | 0.18 ± 0.03 | 2.42 ± 0.09 | 0.81 ± 0.03 | 2.16 ± 0.17 |
| | 25 | 1.01 ± 0.31 | 0.20 ± 0.03 | 2.17 ± 0.09 | 0.79 ± 0.03 | 1.93 ± 0.18 |
| | 30 | 0.88 ± 0.21 | 0.21 ± 0.06 | 1.93 ± 0.07 | 0.78 ± 0.06 | 1.70 ± 0.17 |
| | 36 | 0.79 ± 0.17 | 0.25 ± 0.06 | 1.67 ± 0.04 | 0.74 ± 0.06 | 1.44 ± 0.13 |
| | 41 | 0.77 ± 0.16 | 0.31 ± 0.07 | 1.51 ± 0.03 | 0.68 ± 0.07 | 1.27 ± 0.11 |
| | 46 | 0.73 ± 0.11 | 0.4 ± 0.11 | 1.38 ± 0.04 | 0.6 ± 0.11 | 1.13 ± 0.09 |
| | Temp. [ºC] | $\tau_{MP}$ $[ns]$ | $\alpha_{MP}$ | $\tau_{LP}$ $[ns]$ | $\alpha_{LP}$ | $\tau_{Avg}$ $[ns]$ |
| FGC12 | 10 | 1.66 ± 0.10 | 0.30 ± 0.01 | 3.50 ± 0.07 | 0.69 ± 0.01 | 2.94 ± 0.09 |
| | 15 | 1.70 ± 0.05 | 0.34 ± 0.04 | 3.37 ± 0.05 | 0.65 ± 0.04 | 2.79 ± 0.07 |
| | 20 | 1.67 ± 0.07 | 0.38 ± 0.01 | 3.18 ± 0.06 | 0.61 ± 0.01 | 2.59 ± 0.07 |
| | 25 | 1.51 ± 0.08 | 0.39 ± 0.02 | 2.92 ± 0.05 | 0.60 ± 0.02 | 2.37 ± 0.07 |
| | 30 | 1.41 ± 0.11 | 0.42 ± 0.01 | 2.70 ± 0.06 | 0.57 ± 0.01 | 2.15 ± 0.07 |
| | 36 | 1.28 ± 0.11 | 0.49 ± 0.03 | 2.45 ± 0.04 | 0.51 ± 0.03 | 1.88 ± 0.07 |
| | 41 | 1.16 ± 0.13 | 0.53 ± 0.05 | 2.24 ± 0.09 | 0.46 ± 0.05 | 1.67 ± 0.07 |
| | 46 | 1.03 ± 0.10 | 0.55 ± 0.03 | 2.01 ± 0.07 | 0.45 ± 0.03 | 1.47 ± 0.06 |

Table S3: Values of the fit parameters of fluorescence lifetime histograms across different temperature of MCF-7 derived GPMVs at $\lambda_{em}$ = 560 nm.

| Sample | Temp. [ºC] | $\tau_{MP}$ [ns] | $\alpha_{MP}$ | $\tau_{LP}$ [ns] | $\alpha_{LP}$ | $\tau_{photo}$ [ns] | $\alpha_{photo}$ | $\tau_{Avg}$ [ns] |
|---|---|---|---|---|---|---|---|---|
| | 10 | 1.45 ± 0.05 | 0.3 | 3.75 ± 0.02 | 0.24 | 0.26 ± 0.01 | 0.45 | 1.46 ± 0.06 |

| | | | | | | | | |
|---|---|---|---|---|---|---|---|---|
| GPMVs | 13 | 1.44 ± 0.05 | 0.31 | 3.69 ± 0.02 | 0.23 | 0.26 ± 0.01 | 0.46 | 1.42 ± 0.06 |
| | 16 | 1.36 ± 0.05 | 0.3 | 3.55 ± 0.02 | 0.24 | 0.25 ± 0.01 | 0.45 | 1.38 ± 0.06 |
| | 19 | 1.39 ± 0.05 | 0.31 | 3.52 ± 0.02 | 0.23 | 0.25 ± 0.01 | 0.46 | 1.35 ± 0.06 |
| | 22 | 1.35 ± 0.05 | 0.31 | 3.43 ± 0.02 | 0.23 | 0.25 ± 0.01 | 0.46 | 1.32 ± 0.06 |
| | 25 | 1.37 ± 0.06 | 0.31 | 3.38 ± 0.02 | 0.22 | 0.26 ± 0.01 | 0.47 | 1.29 ± 0.06 |
| | 28 | 1.32 ± 0.06 | 0.31 | 3.27 ± 0.02 | 0.22 | 0.24 ± 0.01 | 0.47 | 1.23 ± 0.06 |
| | 31 | 1.24 ± 0.04 | 0.3 | 3.17 ± 0.01 | 0.21 | 0.20 ± 0.01 | 0.49 | 1.13 ± 0.05 |
| | 34 | 1.20 ± 0.04 | 0.3 | 3.08 ± 0.02 | 0.21 | 0.19 ± 0.01 | 0.49 | 1.09 ± 0.05 |
| | 37 | 1.17 ± 0.04 | 0.3 | 3.00 ± 0.02 | 0.21 | 0.19 ± 0.01 | 0.49 | 1.06 ± 0.05 |
